# The *Drosophila* chromodomain protein Kismet activates steroid hormone receptor transcription to govern axon pruning and memory *in vivo*

**DOI:** 10.1101/376335

**Authors:** Nina K. Latcheva, Jennifer M. Viveiros, Daniel R. Marenda

**Affiliations:** Department of Biology, Drexel University, Philadelphia, PA; Program in Molecular and Cellular Biology and Genetics, Drexel University College of Medicine, Philadelphia, PA; Department of Neurobiology and Anatomy, Drexel University College of Medicine, Philadelphia, PA

**Keywords:** *Drosophila*, *kismet*, axon pruning, Ecdysone Receptor, CHARGE Syndrome, CHD7

## Abstract

Axon pruning is critical for the proper synapse elimination required for sculpting precise neural circuits. Though axon pruning has been described in the literature for decades, relatively little is known about the molecular and cellular mechanisms that govern axon pruning *in vivo*. Here, we show that the epigenetic reader Kismet (Kis) binds to cisregulatory elements of the steroid hormone receptor *Ecdysone Receptor* (*EcR*) gene in *Drosophila* neurons. Kis is required to activate *EcR* transcription at these elements and promote H3K36 di- and tri-methylation and H4K16 acetylation. We show that exogenous EcR can rescue axon pruning and memory defects associated with loss of Kis, and that the histone deacetylase inhibitor SAHA can rescue these phenotypes. EcR protein abundance is a cell-autonomous, rate-limiting step required to initiate axon pruning in *Drosophila,* and our data suggests that this step is under epigenetic control.

**Highlights (bullet points up to 4):**

- The chromodomain reader Kismet activates transcription of the steroid hormone receptor *EcR-B1* in the *Drosophila* to initiate developmental axon pruning.
- Kismet promotes H3K36 di- and tri-methylation and H4K16 acetylation at the *EcR* locus and upstream cis-regulatory sites.
- Axon pruning and memory defects associated with loss of Kismet are significantly rescued by general HDAC inhibition.

## Introduction

The elimination and refinement of synaptic connections is an integral part of normal development in vertebrates and invertebrates alike. Early in the developing nervous system periods of progressive growth result in an overelaboration of synaptic connections onto a target. Inappropriate synapses then need to be eliminated to establish functional organization of the neuronal circuitry (Tau and Peterson, 2010). The pruning of these exuberant connections can occur on a small scale, as with dendritic remodeling, or on a large scale such as with axon retraction and degeneration; with each type occurring through distinct molecular mechanisms (Low and Cheng, 2006). Precise control of axon pruning is critical for proper nervous system function, as defects in pruning have been well documented to lead to developmental neurological and psychiatric disorders (Tau and Peterson, 2010). Despite its vital role, relatively little is known about the mechanisms that govern axon pruning *in vivo*. What is well known about this process is that it requires tight regulation of gene expression in order to execute the necessary signaling pathways in a temporal and tissue specific manner (Awasaki et al., 2011; Yu and Schuldiner, 2014). Epigenetic regulation is key to orchestrating precise gene expression programs for many tightly controlled processes in the body. However, its involvement in axon pruning is still unclear.

Holometabolous insects provide an attractive model for studying axon pruning as their nervous system undergoes extensive reorganization during metamorphosis (Levine et al., 1995; Truman, 1990). In *Drosophila melanogaster* the larval neuronal circuitry is eliminated to make way for adult specific circuitry that governs adult behavior. The most notable changes occur in the learning and memory processing center of the fly brain known as the mushroom bodies (MBs) (Connolly et al., 1996; Ferveur et al., 1995; Heisenberg et al., 1985; McBride et al., 1999). The MBs are bilateral symmetrical structures in the central brain that are comprised of ∼2500 Kenyon cells which belong to 5 types of neurons: gamma, alpha, beta, alpha prime, and beta prime (Ito et al., 1997; Lee et al., 1999). Each Kenyon cell has dendrites which extend into a structure called the calyx, as well as densely packed axons which make up the peduncle. From the peduncle, the axons then divide to form two separate lobes that extend into the dorsal and medial direction. The gamma neurons are generated first in development and initially extend bifurcated axons in the dorsal and medial lobes (Ito et al., 1997; Lee et al., 1999; Spindler and Hartenstein, 2010; Technau and Heisenberg, 1982). During metamorphosis however, the gamma neuron axons are selectively pruned back to the peduncle to eliminate the bifurcation. At approximately 18-22 hours after puparium formation, the gamma neuron axons begin to re-extend new axons only in the medial lobe. This stereotypical developmental pruning of the gamma neurons has been shown to be initiated by the steroid hormone 20-hydroxyecdysone (ecdysone) (Awasaki et al., 2011; Lee et al., 2000; Zheng et al., 2003).

Ecdysone is most well known as the major molting hormone for its role in initiating each of the developmental transitions in arthropods (Handler, 1982). In *Drosophila*, ecdysone is released in large quantities by the prothoracic gland prior to each of the larval molts and pupation. The ligand is then able to enter the cytoplasm of target cells where it can bind to the Ecdysone Receptor (EcR). The binding of ecdysone to EcR stabilizes its interaction with its dimerization partner Ultraspiracle (USP) (Thummel, 2002; Yamanaka et al., 2013). The stable heterodimer is then able to enter the nucleus and activate transcription of a small subset of regulatory target genes known as immediate early genes which possess ecdysone response elements (ERE) in the promotor regions (Ashburner, 1974; Ashburner et al., 1974; Thummel, 2002; Yamanaka et al., 2013). The specific responses different tissues have to induction of the ecdysone signaling cascade can be correlated to the three different types of EcR isoforms expressed in Drosophila: EcR-A, EcR-B1, and EcR-B2 (Talbot et al., 1993; Truman, 1990; Truman et al., 1994). The gamma neurons of the MBs in particular express EcR-B1 which has been shown to be a rate-limiting and cell-autonomous step required for the developmental pruning of axons during metamorphosis (Lee et al., 2000). In addition, EcR-B1 and functional gamma neurons in adult flies were shown to be required for short-term memory and formation of courtship associated long-term memory (Boulanger and Dura, 2015; Ishimoto et al., 2009; Redt-Clouet et al., 2012).

Our lab previously identified the chromodomain protein Kismet (Kis) as necessary for proper developmental axon pruning in the *Drosophila* MB neurons, although the mechanism by which Kis accomplished this was unknown (Melicharek et al., 2010). Kis is the *Drosophila* ortholog of the mammalian chromatin ATPase CHD7, a chromatin “reader” that is thought to play a role in chromatin remodeling by binding to methylated histone tails (Layman et al., 2010). In humans, heterozygous mutations in *CHD7* cause CHARGE syndrome (Vissers et al., 2004), an autosomal dominant neurodevelopmental disorder.

Here, we investigated the role of Kis in the developmental axon pruning of the *Drosophila* MB neurons. We determined that the loss of Kis in the MBs results in pruning defects during metamorphosis, which persist into adulthood and are due to a decrease in transcription of *EcR-B1*. We show that endogenous Kis is enriched at cis-regulatory enhancer sites as well as the transcriptional start site (TSS) of the *EcR* locus. Further, loss of Kis leads to a decrease in the histone marks H3K36me2 and H3K36me3, which have been associated with actively transcribed genes in flies. Additionally, loss of Kis resulted in a striking loss of global H4K16 acetylation. Adult flies with Kis specifically decreased in the MB neurons display a loss of immediate recall memory, which is rescued by exogenous expression of EcR-B1. Finally, we show that pharmacological intervention via the general histone deacetylase (HDAC) inhibitor suberoylanilide hydroxamic acid (SAHA) can rescue the decrease in *EcR-B1* mRNA, axon pruning, and memory defects associated with decreased Kis in MB neurons. Taken together, these data show that Kis-mediated regulation of *EcR-B1* is required for proper developmental axon pruning *in vivo* by mediating the epigenetic marks H3K36me2, H3K36me3, and H4K16ac. These novel findings suggest that the rate-limiting step required to initiate axon pruning (*EcR-B1* expression) is under epigenetic control.

## Results

### Kismet is required for MB pruning

To characterize the pruning defects previously observed in *kis* mutant MB neurons, we utilized the mosaic analysis with a repressible cell marker (MARCM) system to generate homozygous mutant neuroblast clones tagged with a membrane bound GFP using the *201Y*-Gal-4 driver (Lee and Luo, 1999; Melicharek et al., 2010; Schuldiner et al., 2008; Yang et al., 1995). We measured dorsal, medial, and total surface area of the MB lobes in pupal brains 18-22 hours after puparium formation (APF). At this timepoint, the MB lobes are mostly pruned away in control animals (Fig. 1A, 1E). In the null mutant *kis*^*LM27*^, MB clones had significantly larger medial and total lobe surface areas compared to control MB clones (Figs. 1A, 1B, 1E). MB clones expressing the wild-type Kis-L protein in the *kis*^*LM27*^ mutant background showed a significant reduction of the medial and total lobe areas back to control levels (Figs. 1D, 1E), suggesting the defect is due to loss of *kis* function. Overexpression of Kis-L alone did not have an effect on lobe surface area (Figs. 1C, 1E). In addition to MARCM analysis, we utilized RNA interference (RNAi) mediated knockdown of Kis with two separate previously validated RNAi constructs to verify the pruning defects (Melicharek et al., 2010). Pan-neural knockdown of Kis using the *elav*-Gal-4 driver showed a significant increase in the medial and total lobe areas compared to outcross controls (Fig. S1). Similar to the MARCM analysis, pan-neural expression of Kis-L in the knockdown genetic backgrounds was able to significantly rescue the medial and total surface area levels (Fig. S1). Taken together, these data show that Kis is required for the developmental axon pruning of MB neurons during metamorphosis.

**Figure 1.**
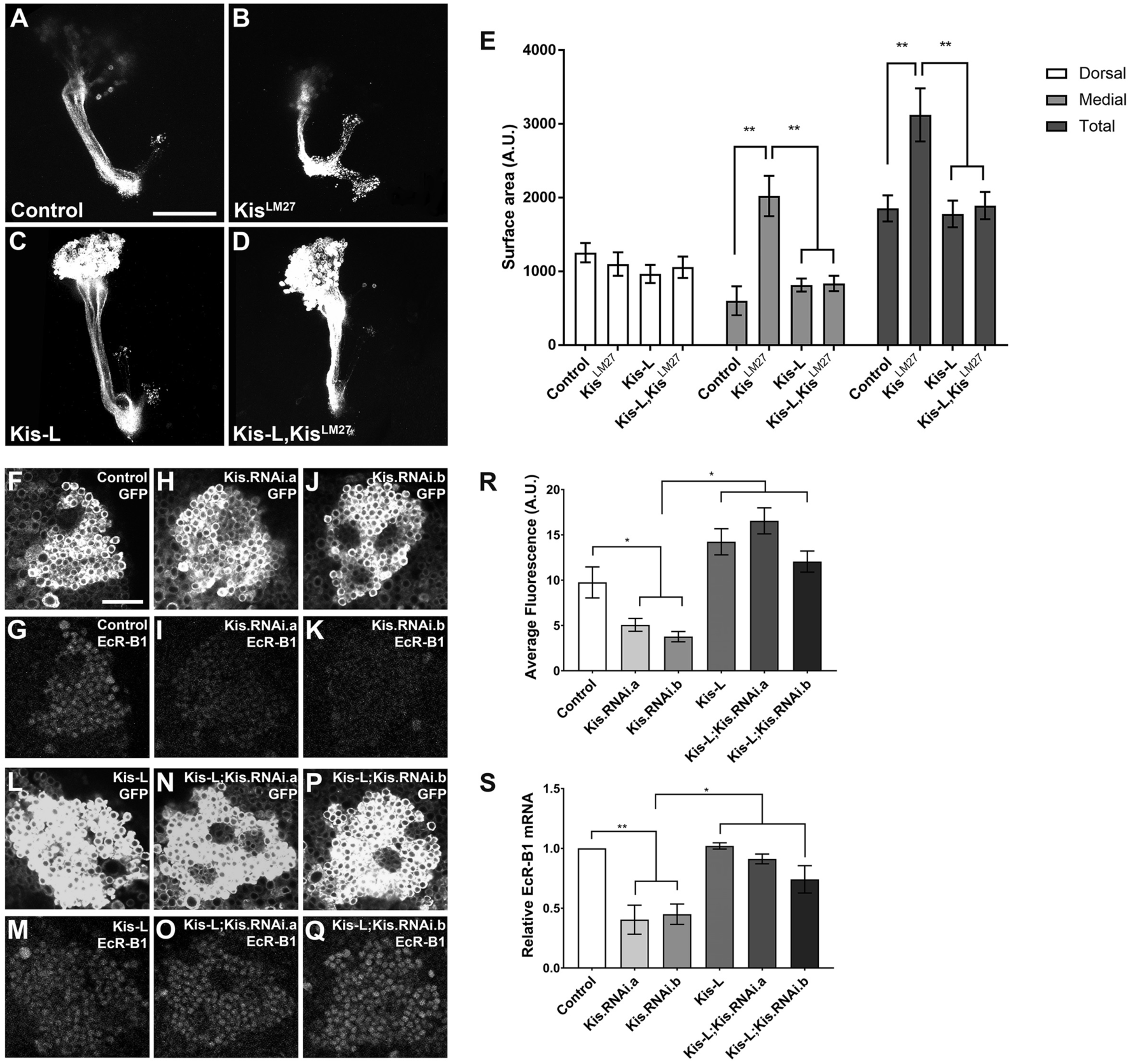
Kis is required for axon pruning and EcR expression. (A-D) Representative images of MARCM generated MB clones expressing membrane bound GFP using the *201Y*-Gal-4 driver 18-22 hours APF. (E) Quantification of dorsal, medial, and total MB lobe surface areas in MARCM animals (n = 8-12). (F-Q) Representative *elav*-Gal-4 driven control, knockdown, overexpression, and rescue pupal brains expressing membrane bound GFP or stained with α-EcR-B1, as indicated. (R) Quantification of fluorescent intensity of EcR-B1 in *elav*-Gal-4 driven pupae (n = 9-12). (S) *EcR-B1* mRNA levels in pupal brains of *elav*-Gal-4 driven animals analyzed by RT-qPCR (n = 3-4). Scale is 10 μm in (A) and 20 μm in (F).

### Kismet binds EcR and activates EcR transcription

Expression of the steroid hormone receptor EcR-B1 is the first step in the developmental axon pruning of the MB neurons, and loss of EcR-B1 function produces defects in pruning similar to what we observe with decreased Kis function (Lee et al., 2000). Therefore, we sought to determine if Kis affects levels of EcR-B1. RNAi knockdown of Kis in MB neurons *in vivo* showed a significant decrease in EcR-B1 immunofluorescence compared to control MBs (Figs. 1F-1K, Fig. 1R). In support of this, mRNA levels of EcR-B1 were significantly reduced in pupae with decreased Kis as well (Fig. 1S). Finally, we wanted to determine if addition of exogenous Kis-L in Kis knockdown MBs could rescue the loss of EcR-B1 expression. We found that replacement of Kis-L protein did successfully rescue both decreased EcR-B1 mRNA and protein expression in these neurons (Figs. 1L-1S). Taken together, these data suggest that Kis is required to promote EcR-B1 expression within MB neurons.

Given that Kis is an epigenetic chromatin reader, we theorized that it may be affecting EcR-B1 mRNA and protein levels by promoting transcription. To test this, we performed chromatin immunoprecipitation (ChIP) followed by qPCR to analyze Kis occupancy at previously published cis-regulatory elements of the *EcR* locus as well as the *EcR* TSS (Fig. 2A) (Boulanger et al., 2011). We utilized a Kis-eGFP protein trap animal previously described to express an eGFP tagged version of the Kis protein (Buszczak et al., 2007; Ghosh et al., 2014). Importantly, this animal shows normal localization and function of Kis, and we have shown that our RNAi reagents can successfully knockdown the eGFP tagged Kis protein and *kis* mRNA (Fig. S2) (Ghosh et al., 2014). We utilized the *forkhead* (*fkh*) TSS as a positive control, as it was previously reported to be bound by endogenous Kis (Srinivasan et al., 2008a). Additionally, we used the *dynamin* homolog *shibire* (*shi*) as a negative control, as we had previously shown via microarray analysis that loss of Kis did not have any significant effect on *shi* expression (Ghosh et al., 2014). We verified that Kis was not enriched at the *shi* promoter region (Fig. 2A), and that knockdown of Kis did not have any significant effect on Kis abundance at the *shi* promoter (Fig. 2A). In contrast, wild-type control brains showed enrichment of Kis at the two previously reported *EcR* cis-regulatory elements (*EcR.1* and *EcR.2*) as well as the *EcR* and *fkh* TSS (Fig. 2A). Finally, we observed a significant decrease in Kis enrichment at the *fkh* TSS, *EcR.1, EcR.2*, and *EcR* TSS sites when Kis was knocked down, confirming specificity of Kis binding at these sites (Fig. 2A). These results suggest that Kis binds to known cis-regulatory sites and the TSS of the *EcR* locus in *Drosophila* larval brains.

**Figure 2.**
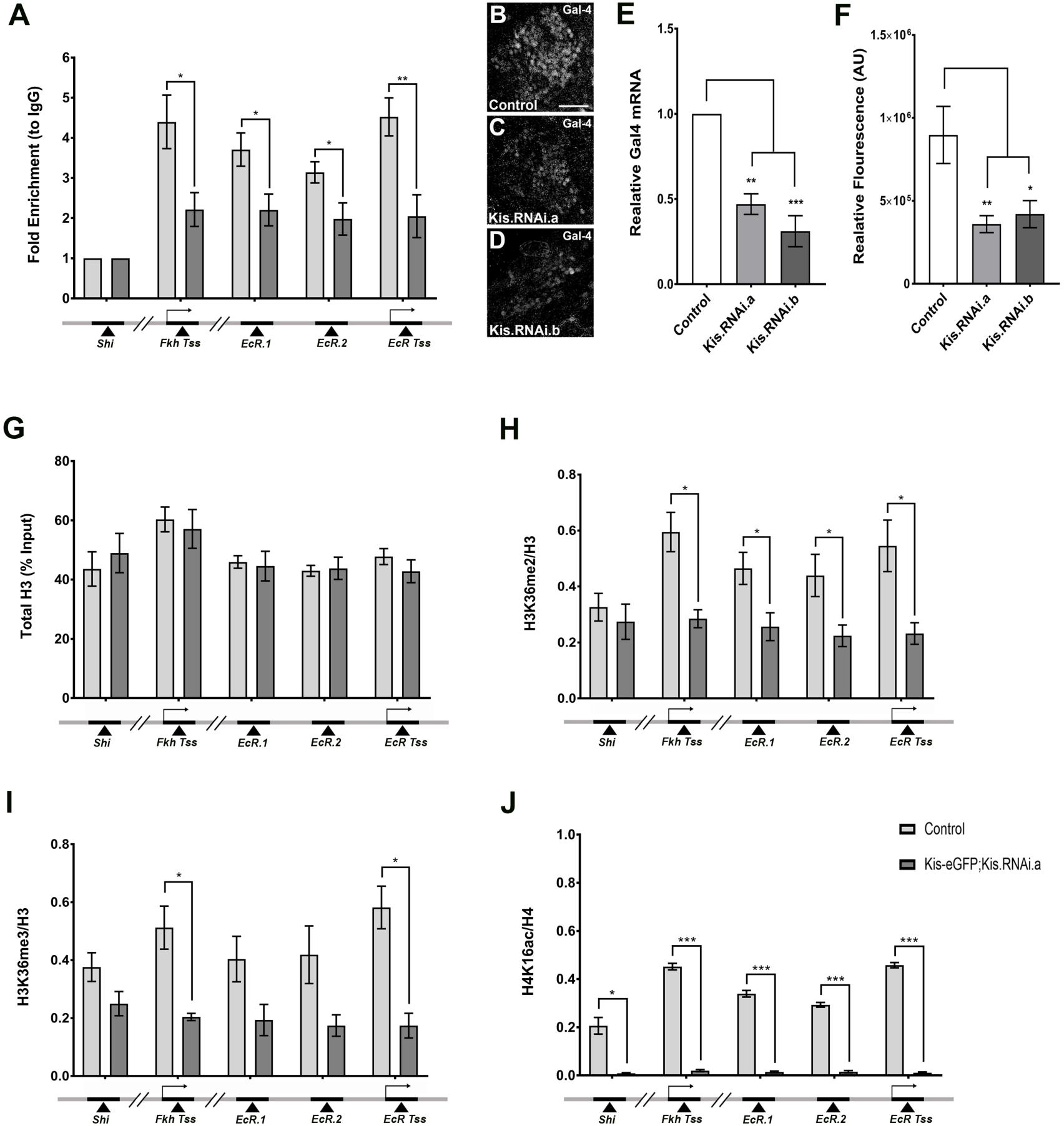
Kis binds to EcR loci promoting EcR-B1 transcription and H3K36 methylation. (A) ChIP-qPCR analysis using chromatin isolated from 3^rd^ instar larval brains. Kis enrichment at the *EcR* TSS and enhancer sites, the *fkh* TSS, and the *shi* promoter site (n = 4-5). (B-D) Representative images of control and knockdown *EcR.1*-Gal-4 driven pupal brains stained with α-Gal-4. (E) Quantification of fluorescent intensity of α-Gal-4 (n = 10). (F) *Gal-4* mRNA levels of *EcR.1*-Gal-4 driven animals analyzed by RT-qPCR (n = 3). (G) Total H3 as a percentage of the input DNA was determined by qPCR at the previously noted genomic loci (n = 6). (H, I) ChIP-qPCR analysis of H3K36me2 and H3K36me3 abundance relative to total H3 at the above mentioned genomic loci, respectively (n = 3). (J) ChIP-qPCR analysis of H4K16ac abundance relative to total H4 at the above mentioned genomic loci, respectively (n = 3). Scale is 20 μm in (B).

To determine if Kis can control transcriptional output from these sites, we utilized an *EcR* transcriptional reporter (*EcR.1*-Gal-4, described in (Pfeiffer et al., 2011; Pfeiffer et al., 2008)). This reporter drives the expression of the exogenous Gal-4 protein from the *EcR.1* cis-regulatory element that is partially responsible for the expression of *EcR-B1* in MB neurons (Boulanger et al., 2011). Using this reporter, we determined the levels of Gal-4 mRNA and protein in wild-type control MBs and Kis knockdown MBs. We observed a significant decrease in both Gal-4 mRNA and protein levels from this reporter in Kis knockdown animals compared to controls (Figs. 2C-G). Taken together, these data suggest that Kis protein can bind to *EcR* cis-regulatory elements and the *EcR* TSS to promote transcriptional activation and increase EcR-B1 expression in the *Drosophila* nervous system.

### Kismet promotes H3K36 methylation and H4K16 acetylation

One mechanism by which chromatin remodeling proteins promote transcriptional activation is by mobilizing nucleosomes and allowing transcriptional machinery access to enhancer sites and promoters of target genes (Clapier et al., 2017). Given Kis’s homology to CHD7 and its conserved ATPase domain, we hypothesized that Kis may be removing nucleosomes from the *EcR* locus to allow access to the transcriptional machinery. To test this possibility, we immunoprecipitated total histone 3 (H3) protein at the *EcR* locus in control and Kis knockdown brains. However, we found no significant difference between these conditions at each of the cis-regulatory loci we analyzed (Fig. 2B). To verify this, we performed an MNase protection assay. Consistent with our total H3 analysis, we saw no change in the digestion of chromatin upon Kis knockdown (Fig. S3). Taken together, these data suggest that Kis is not affecting the movement of nucleosomes as a mechanism to promote transcription at the *EcR* locus.

Another way chromatin readers can affect gene expression is by altering the histone modifications present at relevant genomic loci (Clapier et al., 2017), and Kis has been shown to affect histone modifications previously (Srinivasan et al., 2008a). We began by analyzing H3K4 methylation states at *EcR* loci in control and Kis knockdown animals, as it is the type of methylation most commonly associated with actively transcribed genes. ChIP-qPCR analysis showed no change in all types of H3K4 methylation levels upon Kis knockdown at these loci (Figs. S4A-C). We next analyzed H3K27 trimethylation, as this modification is often associated with transcriptional repression and previously shown to be increased in Kis mutant salivary glands (Srinivasan et al., 2008b). We did not observe any significant change in H3K27 trimethylation upon Kis knockdown at any of the loci examined (Fig. S4D). Taken together, these data suggest that alteration of H3K4 methylation or H3K27 trimethylation is not part of the mechanism by which Kis controls *EcR* transcription in the *Drosophila* CNS.

Previous studies demonstrated a global decrease in H3K36 di- and tri-methylation upon Kis loss in *Drosophila* salivary gland polytene chromosomes (Dorighi and Tamkun, 2013). This type of modification is usually associated with actively transcribed genes in *Drosophila* (Dorighi and Tamkun, 2013; Stabell et al., 2007; Wagner and Carpenter, 2012). We therefore sought to determine if the same effect on H3K36 methylation was present in the *Drosophila* nervous system upon pan-neural knockdown of Kis protein. We observed that H3K36me2 was significantly decreased at all the *EcR* cis-regulatory sites, as well as at the *fkh* positive control, in Kis knockdown brains compared to controls (Fig. 2H). Additionally, H3K36me3 was also significantly decreased at the *EcR* and *fkh* TSS (Fig. 2I). Importantly, this decrease in H3K36 methylation was not global, as there was no significant change with either H3K36 di- or tri-methylation at the *shi* promoter which is not bound by Kis (Figs. 2A, 2H-I). Also, given that H3K36me2 is the substrate for the trimethylated form, it is likely that the decrease in H3K36me3 is due to the decrease in H3K36me2. Combined, these results suggest Kis affects transcription at the *EcR* locus by promoting the active H3K36 di- and tri-methylation histone marks.

Previous studies with H3K36me2 have demonstrated a synergistic relationship with other histone modifications, particularly H4K16 acetylation (H4K16ac) (Bell et al., 2007). H4K16 acetylation directly influences transcription by positively regulating chromatin accessibility to non-histone proteins (Zhang et al., 2017). Additionally, this type of post-translational modification has been shown to negatively impact chromatin condensation by preventing the function of ATP-dependent chromatin-assembly factor (ACF) which is involved in the condensation of 30-nanometer chromatin fibers (Shogren-Knaak et al., 2006; Yang et al., 2006). Since we saw a significant decrease in H3K36me2 at our loci of interest, we wanted to determine if H4K16ac levels were also affected in Kis knockdown animals. ChIP-qPCR revealed that H4K16ac was significantly decreased when Kis was knocked down (Fig. 2J) compared to controls. This was consistent at all the loci including *shi* promoter, potentially indicative of a universal decrease in H4K16ac.

### EcR-B1 rescues axon pruning defects

EcR-B1 is well documented to be a key player in initiating the axon pruning cascade in MB neurons (Lee et al., 2000; Yu and Schuldiner, 2014). Given that we have shown Kis acts to promote transcription of *EcR-B1* in MB neurons, we sought to determine if we could rescue the pruning defects we observe is Kis mutants by expression of exogenous EcR-B1. Utilizing the MARCM system, we expressed exogenous EcR-B1 protein within *kis*^*LM27*^ mutant MB clones (Fig. 3). We observed that exogenous expression of EcR-B1 significantly reduced the abnormal pruning observed in *kis*^*LM27*^ mutant MB clones (Figs. 3B, 3D, 3E). We also observed a significant reduction in pruning defects when exogenous EcR-B1 was pan-neurally co-expressed with Kis RNAi constructs (Fig. S5). Interestingly, EcR-B1 overexpression alone, as well as EcR-B1 expression in Kis loss of function backgrounds, produced smaller surface areas in the dorsal and total lobes compared to outcross controls (Figs. 3A, 3C, 3E-3F). This is consistent with Kis functioning upstream of EcR-B1 in the pruning process, as well as with EcR-B1 being the rate-limiting factor for pruning. Taken together, these data suggest that Kis mediates axon pruning in MB neurons by transcriptionally activating *EcR-B1*, thereby controlling EcR-B1 protein levels.

**Figure 3.**
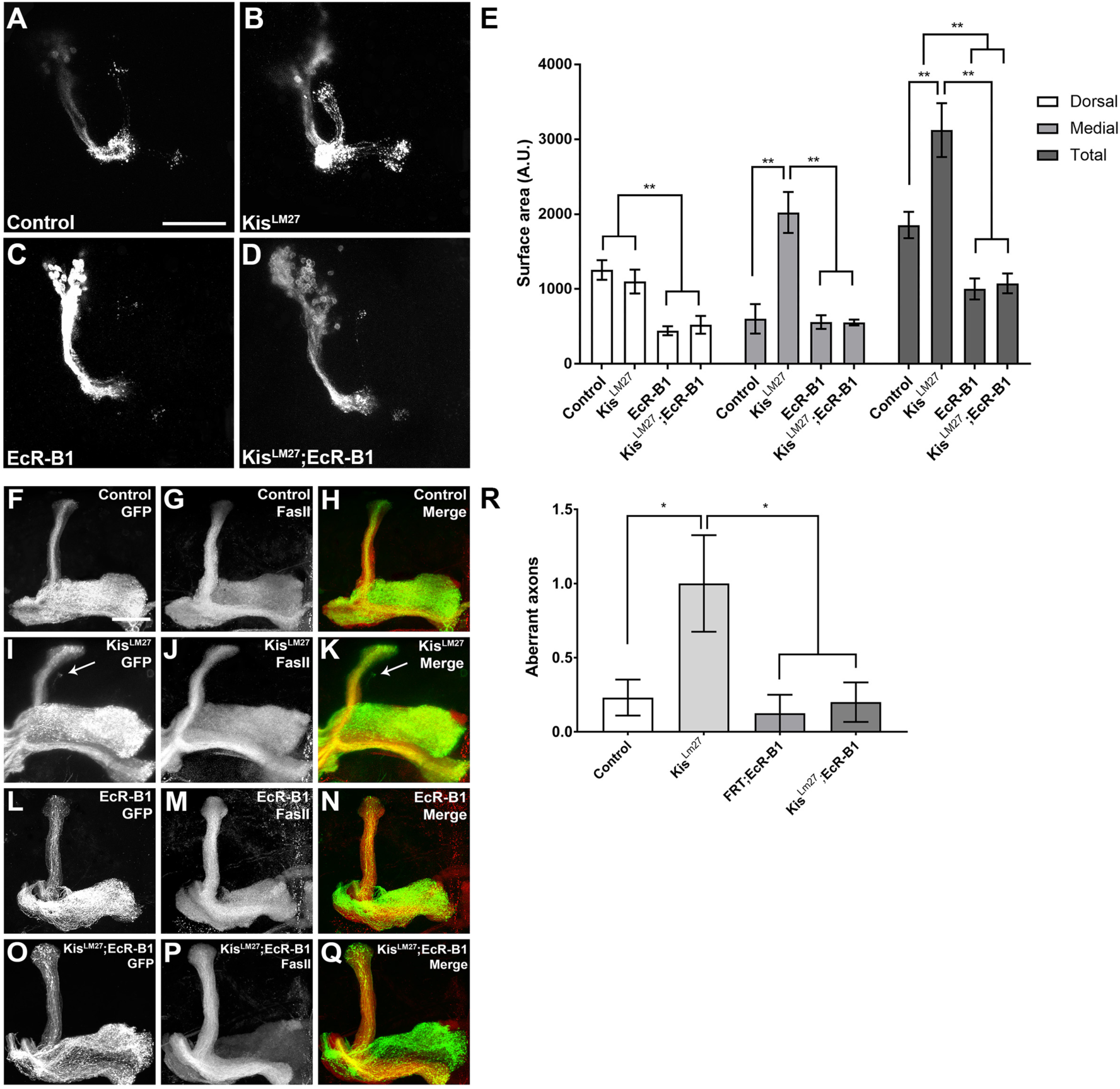
Exogenous EcR-B1 rescues defective axon pruning associated with loss of Kis. (A-D) Representative images of MARCM generated MB clones expressing membrane bound GFP using the *201Y*-Gal-4 driver 18-22 hours APF. (E) Quantification of dorsal, medial, and total MB lobe surface areas in MARCM animals (n = 10-12). (F-Q) Representative adult MARCM generated MB clones expressing membrane bound GFP or stained with α-FasII, as indicated. (R) Quantification of aberrant axons in MBs (n = 10). Scale is 10 μm in (A) and 20 μm in (F).

Pruning defects observed during metamorphosis may simply reflect a delay in normal pruning. Therefore, to determine that homozygous MARCM *kis*^*LM27*^ mutant clones indeed have defective developmental pruning, as opposed to having delayed pruning, we sought to determine if the unpruned axons persisted into adulthood. We generated MARCM clones with homozygous *kis*^*LM27*^ MBs as previously described and aged the adults for 5 days after eclosion. We then immunostained the adult brains with anti-FASII, a transmembrane cell adhesion protein, which is differentially expressed in the separate populations of MB neurons (Bornstein et al., 2015; Stewart and McLean, 2004). FASII expression is lowest in the early born gamma neurons and highest in the late born alpha/beta neurons; therefore, the appearance of GFP labeled MARCM axons in the dorsal lobe that are weakly or unstained for FASII would constitute aberrant unpruned axons that persisted into adulthood (Bornstein et al., 2015). Compared to control MBs, *kis*^*LM27*^ MARCM clones had significantly more weakly stained and/or unstained FASII GFP-positive axons outside the dorsal lobe bundle, indicating that pruning is in fact prevented and not delayed in this mutant (Figs. 3F-3K, 3R). Given that exogenous expression of EcR-B1 in the *kis*^*LM27*^ MARCM background rescued the pruning defects at the pupal stage, we tested whether the rescue effects of exogenous EcR-B1 provided at the pupal stage also persisted into adulthood. We observed that expression of EcR-B1 in *kis*^*LM27*^ mutant MARCM clones showed significantly fewer GFP-positive axons outside the dorsal lobe in adult MB axons, while expression of EcR-B1 alone had no effect (Figs. 3L-3Q, 3R). Collectively, these data suggest that the defective pruning observed during metamorphosis in *kis* loss-of-function MB neurons persists into adulthood.

### EcR-B1 rescues memory defects

Previous studies from our lab showed that reduction of Kis levels in MB neurons produced significant defects in immediate recall memory (Melicharek et al., 2010). Since loss of Kis leads to pruning defects that persist into adulthood, and exogenous EcR-B1 expression is able to rescue the defects, we were interested in determining if exogenous expression of EcR-B1 could also rescue the memory defect associated with loss of Kis function. We utilized the conditioned courtship suppression assay which takes advantage of the innate courting behaviors carried out by male *Drosophila* in response to multimodal signals transduced by females (McBride et al., 2005; Siegel and Hall, 1979; Siwicki et al., 2005). Wild-type males will decrease their rate of courting during training when exposed to an unresponsive female, and continue to court at lower rates even with subsequent receptive females for an average of 1-3 hours after exposure (McBride et al., 2005; Siegel and Hall, 1979; Siwicki et al., 2005). We utilized RNAi to knockdown Kis in MB neurons using the *OK107*-Gal4 driver. Importantly, expression of the Gal4 alone, or Gal4-mediated expression of exogenous EcR-B1 did not produce any learning (Fig. 4A) or memory (Fig. 4B) defects. We observed that males with decreased Kis protein displayed intact learning, as evident by the significant decrease in courtship from the initial to final stages of exposure to an unresponsive female (Fig. 4C). Similarly, males with both decreased Kis and exogenously expressed EcR-B1 also had intact learning (Fig. 4C). However, males with decreased Kis showed abnormal memory (Fig. 4D), as these males had rates of courtship that were not significantly different than sham males, which did not receive exposure to an unreceptive female. In contrast, males with both decreased Kis and exogenously expressed EcR-B1 showed significantly reduced courtship in trained males compared to sham males, indicative of intact immediate recall memory (Fig. 4D). Taken together, these data suggest that the memory defects associated with loss of Kis in the MB neurons are due to decreased EcR-B1 levels.

**Figure 4.**
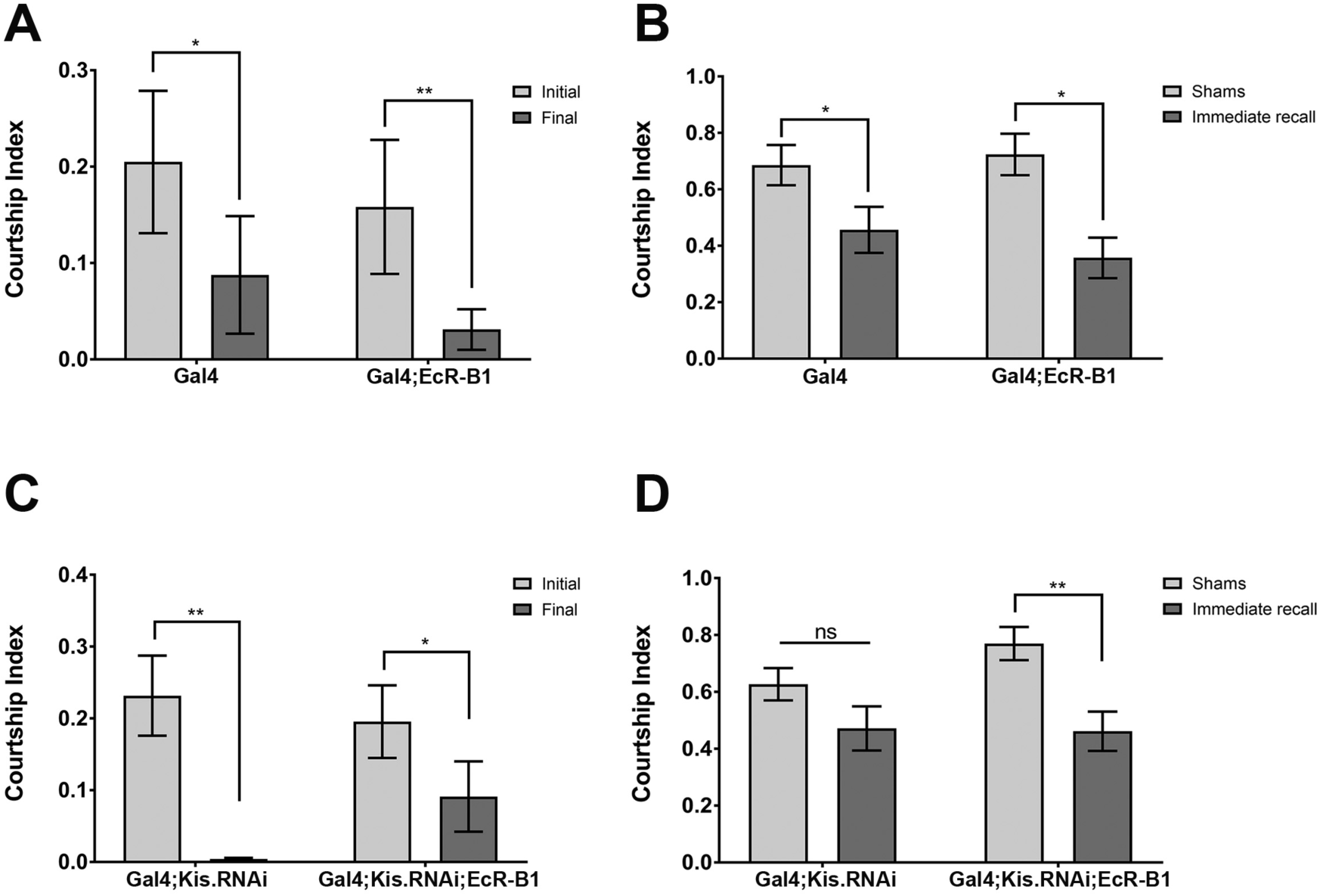
Exogenous EcR-B1 expression rescues memory defects rescued in Kis knockdown MBs. (A, C) Change in courtship index of *OK107*-Gal-4 driven males during the initial (light grey) and final (dark grey) 10 minutes was assessed (n = 20-25). (B, D) Immediate recall of trained *OK107*-Gal-4 driven males was assessed and compared to genetically identical sham trained males (n = 20-25).

### SAHA rescues Kis defects

Recently, our lab showed that pharmacological inhibition of HDACs can rescue multiple defects associated with loss of Kis at the neuromuscular junction (Latcheva et al., 2018). HDAC inhibition (HDACi) did not significantly affect *kis* mRNA levels (Latcheva et al., 2018), so we hypothesized that the rescue may be due to effects on target genes. To further examine this hypothesis, we tested whether HDACi treatment could rescue the decreased EcR-B1 levels, pruning, and memory defects observed in Kis knockdown. We observed that treatment of Kis knockdown animals with SAHA significantly increased *EcR-B1* mRNA levels compared to dimethyl sulfoxide (DMSO) treated controls (Fig. 5S). SAHA treatment alone had no significant impact on axon pruning in either pupal or adult MBs (Figs. 5E, 5R). However, we observed that SAHA treatment significantly decreased the number of unpruned axons in both pupal and adult brains in Kis loss-of-function MB neurons (Figs. 5E, 5R). Finally, SAHA treatment itself did not have an adverse effect on learning or memory in controls (Figs. 6A-6B) and was able to significantly rescue the immediate recall defect in Kis knockdown animals compared to DMSO treatment alone (Figs. 6C-6D). Therefore, SAHA treatment might be counteracting the loss of global H4K16 acetylation we observed in Kis knockdown animals. Taken together, these results show that HDACi treatment significantly rescues multiple defects associated with Kis loss-of-function.

**Figure 5.**
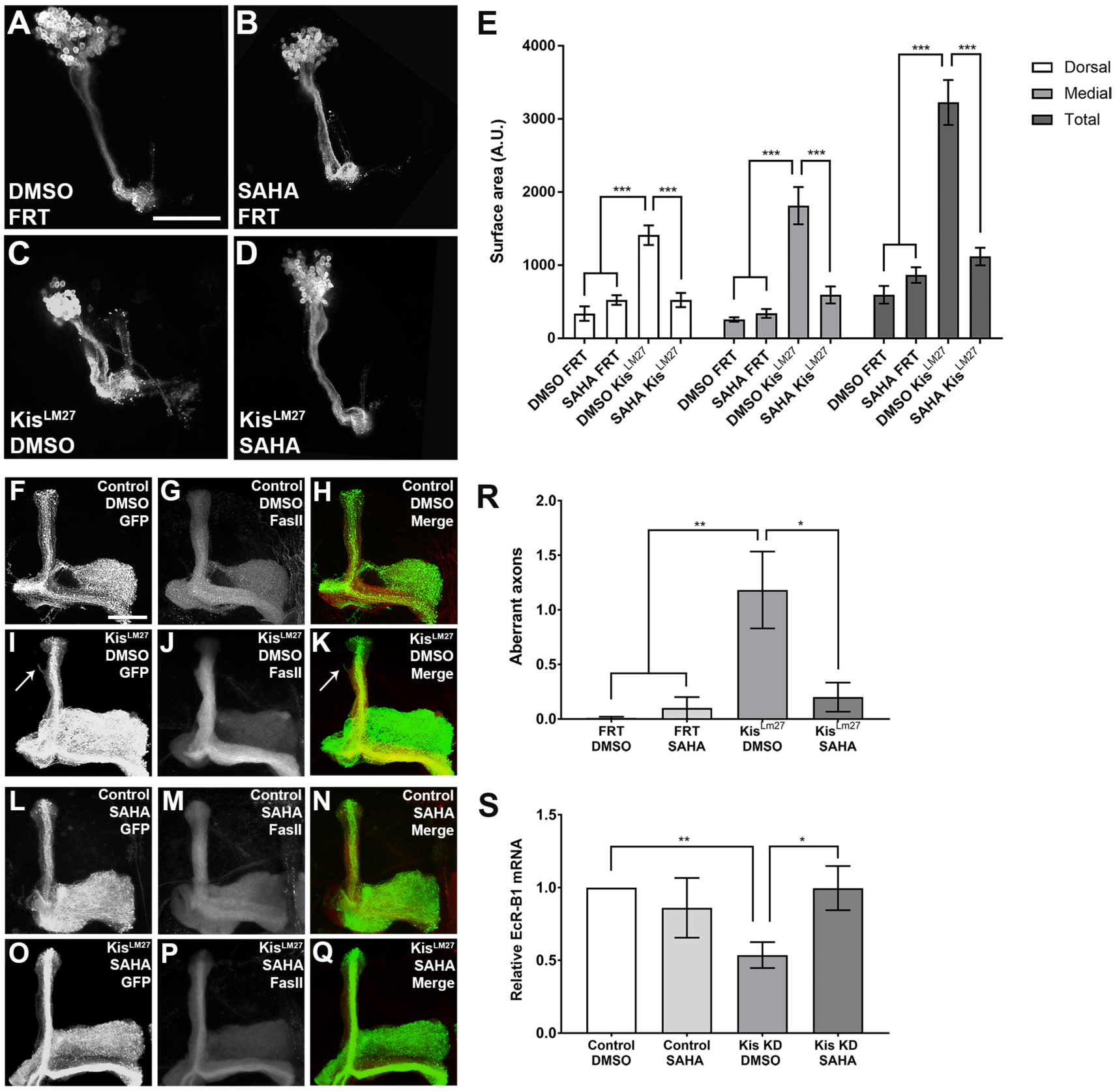
HDAC inhibition rescues defective axon pruning and EcR-B1 expression in loss of Kis animals. (A-D) Representative images of DMSO or SAHA treated MARCM generated MB clones expressing GFP using the *201Y*-Gal-4 driver 18-22 hours APF. (E) Quantification of dorsal, medial, and total MB lobe surface areas in DMSO or SAHA treated MARCM animals (n = 10-12). (F-Q) Representative DMSO or SAHA treated adult MARCM generated MB clones expressing membrane bound GFP or stained with α-FasII, as indicated. (R) Quantification of aberrant axons in MBs of flies treated with DMSO or SAHA (n = 10-12). (S) *EcR-B1* mRNA levels in brains of DMSO or SAHA treated *elav*-Gal-4 driven pupae analyzed by RT-qPCR (n = 3-4). Scale is 10 μm in (A) and 20 μm in (F).

**Figure 6.**
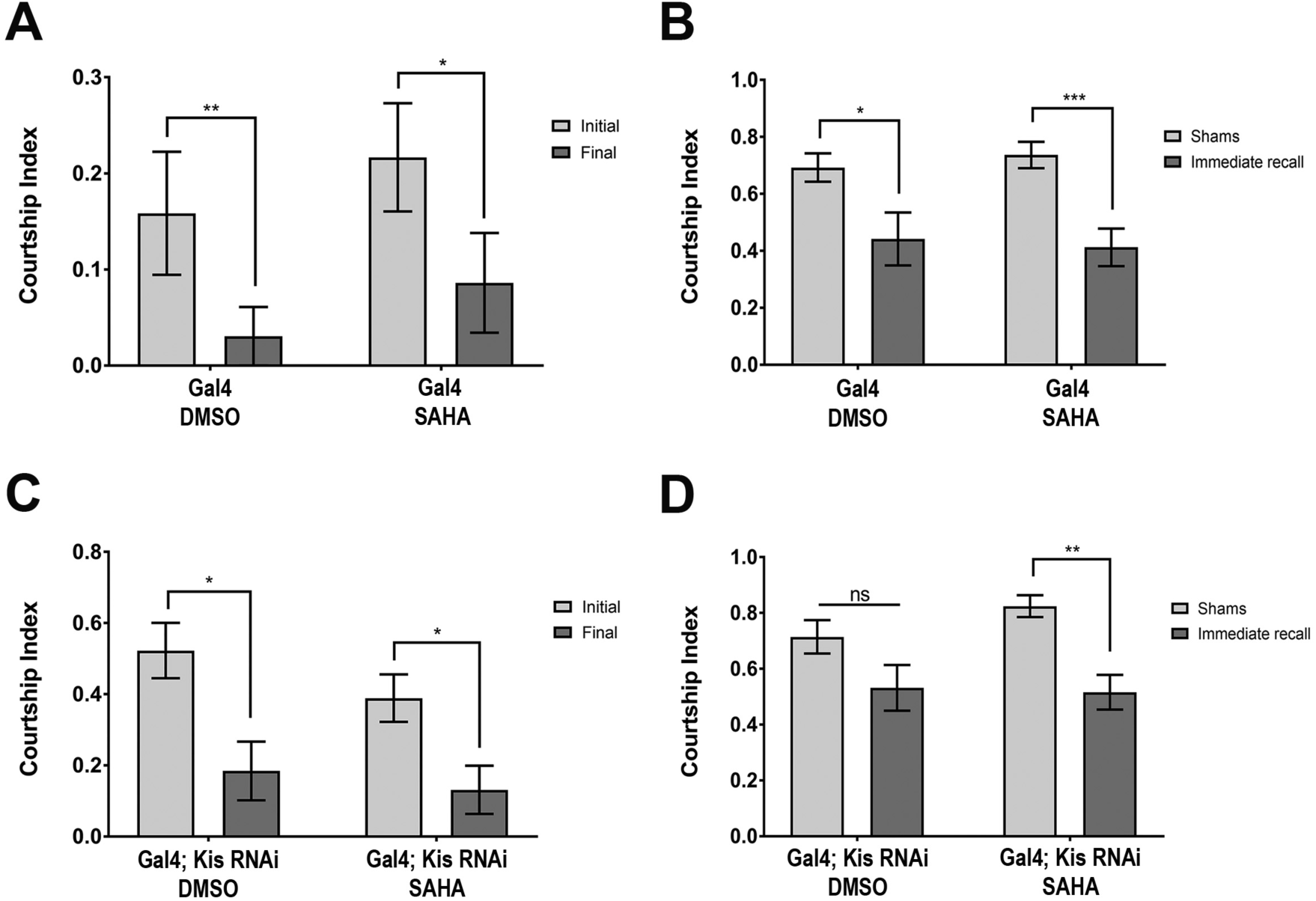
SAHA treatment rescues immediate recall defects rescued by in Kis knockdown animals. (A, C) Change in courtship index of DMSO or SAHA treated *OK107*-Gal-4 driven males during the initial (light grey) and final (dark grey) 10 minutes was assessed (n = 20-25). (B, D) Immediate recall of DMSO or SAHA treated trained *OK107*-Gal-4 driven males assessed and compared to genetically identical and treated sham trained males (n = 20-25).

**Figure 7.**
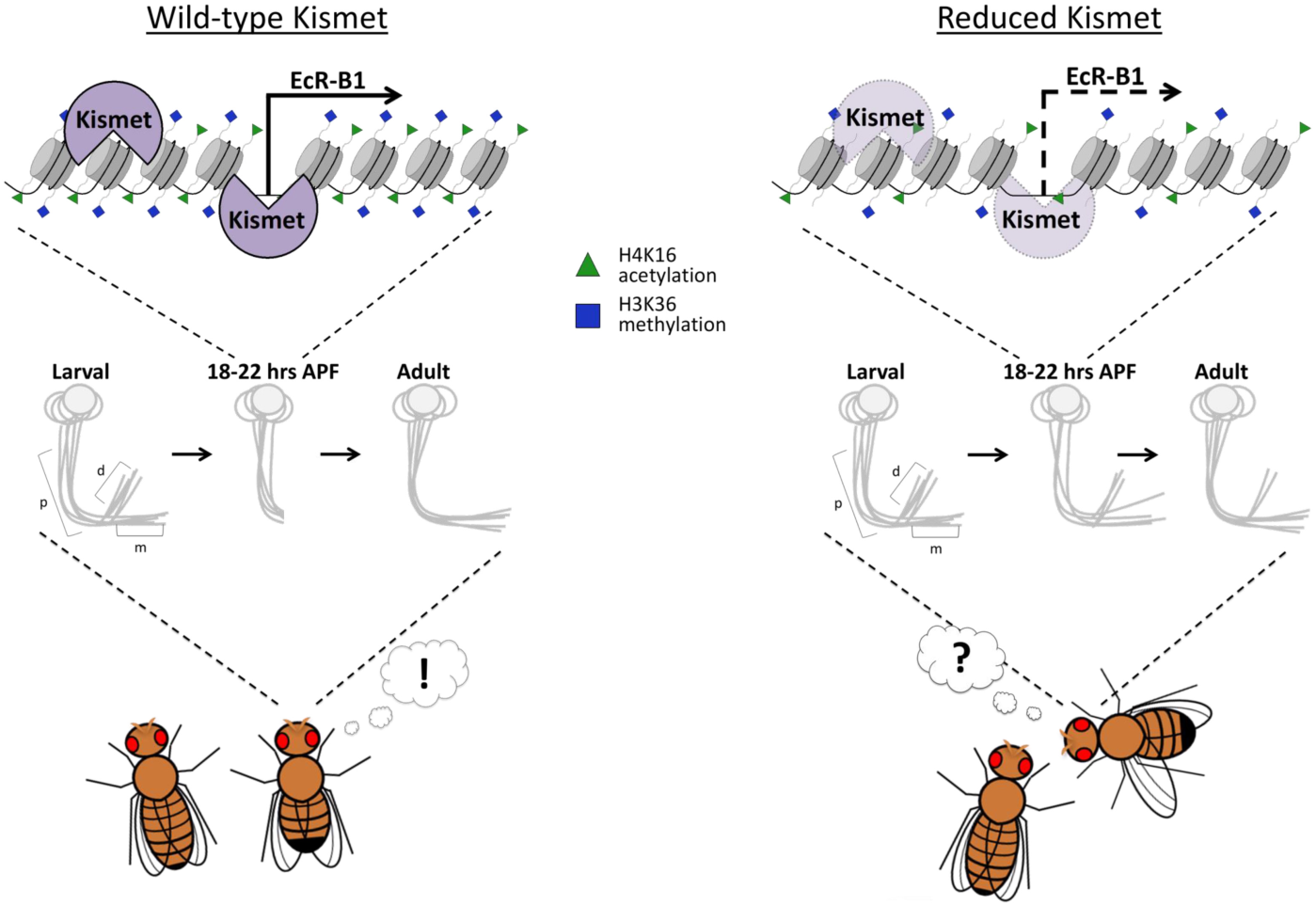
Model of Kis mediated expression of EcR-B1 necessary for MB pruning and behavior. In wild-type flies, Kis promotes the active H3K36 methylation (blue square) and H4K16 acetylation (green triangle) histone marks and binds to cis-regulatory elements and TSS of the *EcR-B1* locus promoting transcription. Kis activates EcR-B1 expression, which is required for proper developmental MB axon pruning, which in turn is necessary for memory formation. In Kis knockdown animals, Kis binding to the *EcR-B1* locus is reduced thereby decreasing H3K36 methylation and H4K16 acetylation marks leading to decreased *EcR-B1* mRNA and protein. Reduction of EcR-B1 leads to defects in MB pruning and immediate recall memory.

## Discussion

Axon pruning and elimination are critical steps to establishing and refining neural circuitry. However, relatively little is known about how the precise extrinsic and intrinsic signals come together to initiate the pruning cascade. Here we begin to unravel the epigenetic mechanisms essential for initiating developmental axon pruning *in vivo*. We show that the chromatin reader Kis activates transcription of *EcR-B1* in the *Drosophila*

MBs and promotes methylation of H3K36 at cis-regulatory sites and the TSS of *EcR-B1*. This loss of H3K36 methylation at specific loci may lead to a global decrease in H4K16 acetylation we observe in Kis knockdown flies. Proper regulation of EcR-B1 by Kis is required for initiating developmental pruning and immediate recall memory in adults. Finally, we show that the general HDACi SAHA can increase *EcR-B1* mRNA levels in animals with decreased Kis and rescue their pruning and memory defects. The SAHA may be acting to reestablish a balance of gene expression due to global loss of H4K16 acetylation when Kis is not present. Taken together, our data shows that the essential rate-limiting step in developmental axon pruning, EcR-B1 expression, is under epigenetic control.

The expression of EcR in the MBs has been subject to studies indicating two distinct pathways (Boulanger et al., 2011; Boulanger and Dura, 2015; Zheng et al., 2003). First, the TGF-β signaling pathway has been implicated in MB gamma neuron remodeling, as mutations in both the Drosophila TGF-β receptor Baboon and its downstream effector dSmad2 both produce pruning defects at 18-22 hours APF (Zheng et al., 2003). Additionally, exogenous expression of EcR-B1 was able to rescue the pruning defects in the TGF-β pathway mutant backgrounds indicating that EcR-B1 is downstream of TGF-β signaling (Zheng et al., 2003). Similarly, our results demonstrate that exogenous EcR-B1 expression bypassed the need for Kis in a mutant background and rescued the pruning defects in both pupal and adult MBs (Fig. 3). This parallels the previous findings by demonstrating that Kis is upstream of EcR-B1 transcription. Where Kis fits relative to TGF-β signaling however still remains to be answered.

A second pathway of activating EcR-B1 in the MBs is via the nuclear receptor Ftz-f1 and its homologue Hr39 has been described (Boulanger et al., 2011; Boulanger and Dura, 2015). Here Ftz-f1 binds directly the EcR cis-regulatory sites and activates EcR-B1 expression while Hr39 acts to block EcR expression in the MBs (Boulanger et al., 2011). This pathway also functions upstream of EcR-B1, however it is independent of TGF-β signaling as Ftz-f1 overexpression did not rescue pruning defects in Baboon mutants (Boulanger et al., 2011). Our data indicates that that Kis is also enriched at the same cis-regulatory sites that Ftz-f1 is found (Fig. 2) but whether they are act in coordination or independently of each other remains to be determined. It may be that both Kis and Ftz-f1 act independently and help create a redundancy in case of the failure of one pathway. Given that reduction of neither Kis nor Ftz-f1 completely eliminated EcR-B1 levels may support this redundancy hypothesis, however more work is necessary to explore the relationship (Boulanger et al., 2011).

Few studies have hinted at an epigenetic mechanism of EcR activation. The *Drosophila* Set2 K36 histone methyl transferase (HMT) was shown to genetically interact with the EcR signaling pathway and positively regulate expression of EcR target genes (Stabell et al., 2007). However, as dSet2 is required for general transcriptional elongation, this may be a global effect of dSet2 function. Nonetheless, this implicates H3K36 methylation as an important factor in EcR expression. While Kis does not appear to affect expression of dSet2, Ash1 or dMes-4 (data not shown), it might be acting to recruit the HMTs to relevant target genes. To this effect, Kis was previously shown to increase the global association of Ash1 on polytene chromosomes (Dorighi and Tamkun, 2013; Srinivasan et al., 2008b). While other studies with Kis have implied a role for the chromatin reader in transcriptional elongation (Srinivasan et al., 2005), we demonstrate a selectivity for Kis binding and Kis-mediated elevation of H3K36 methylation as no significant changes were observed at the *shi* promoter in Kis knockdown animals. Further work needs to be done to tease apart the relationship between Kis, H3K36 methylation, and any HMTs responsible for this modification in the context of EcR regulation.

In the field of epigenetics, much work has been done to understand the crosstalk between different histone modifications and their cumulative outcome on transcription. For example, H3K36me2 was reported to specifically increase H4K16 acetylation, a modification most well-known for its role in de-condensation of chromatin structure and maintenance of active gene expression (Bell et al., 2007; Shogren-Knaak et al., 2006; Zhang et al., 2017). We provide evidence for this type of epigenetic crosstalk by demonstrating a targeted loss of H3K36 methylation and global loss of H4K16 acetylation in Kis knockdown background. It is therefore plausible that Kis might be affecting transcriptional output by promoting H4K16 acetylation via H3K36me2 crosstalk. This may explain why HDAC inhibition with SAHA, seemingly unrelated to histone methylation, is able to restore *EcR-B1* mRNA levels and rescue pruning and memory defects. Furthermore, SAHA treatment has been previously shown to increase H4K16ac and ultimately increase transcriptional output in cancer cells (Barbetti et al., 2013). Additional work is essential to building a complete understanding of the integration of extrinsic and intrinsic cues that control transcriptional regulation. This is fundamental to unraveling the mechanisms that underlie the refinement of neural circuits.

## Acknowledgements

We would like to thank the members of the Marenda lab for their helpful feedback during the course of this project, the Bloomington Drosophila Stock Center, the Vienna Drosophila Stock Center, and the Developmental Studies Hybridoma Bank for fly stocks, antibodies, and reagents; Drexel CIC for imaging analysis and assistance. We would like to thank J. Tamkun, L. Luo, and A. Spralding for stocks. This work was supported by grants from the CHARGE syndrome Foundation (to D.R.M.) and the National Science Foundation IOS 1256114 (to D.R.M.). Some of this material is based upon work supported by (while serving at) the National Science Foundation. Any opinion, findings, and conclusions or recommendations expressed in this material are those of the author(s) and do not necessarily reflect the views of the National Science Foundation.

## Author Contributions

Conceptualization, N.K.L, J.M.V., D.R.M.; Methodology, N.K.L, J.M.V., D.R.M.; Validation, N.K.L, J.M.V.; Formal Analysis, N.K.L, J.M.V., D.R.M.; Investigation, N.K.L, J.M.V.; Writing – Original Draft, N.K.L, J.M.V., D.R.M.; Writing – Review & Editing, N.K.L, J.M.V., D.R.M.; Supervision, D.R.M.; Project Administration, D.R.M.; Funding Acquisition, D.R.M.

---

**Figure S1.**
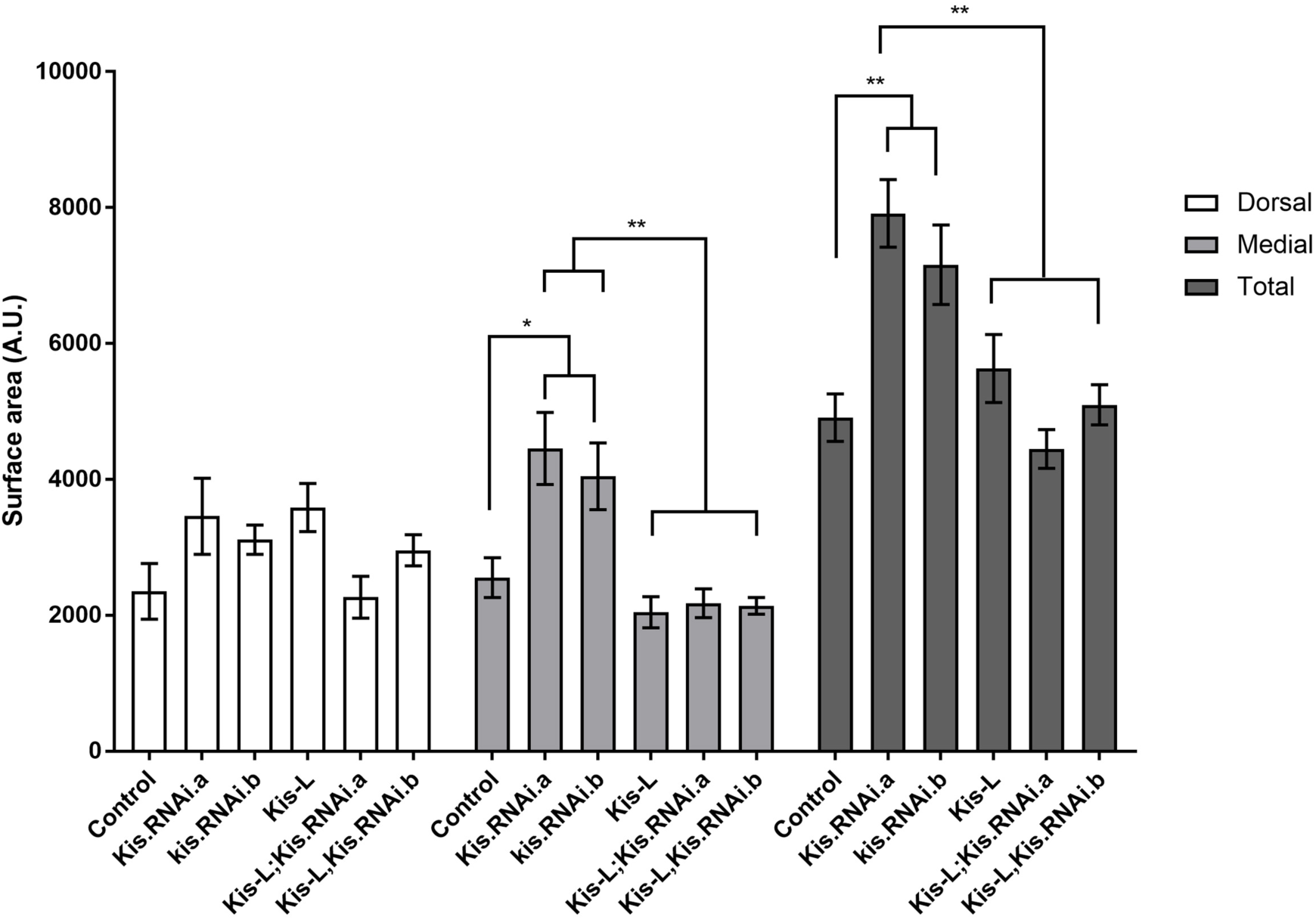
Kis is required for axon pruning in Kis knockdown pupae. Quantification of dorsal, medial, and total MB lobe surface areas in *elav*-Gal-4, mCD8-GFP driven pupal brains stained with α-EcR-B1 (n = 10).

**Figure S2.**
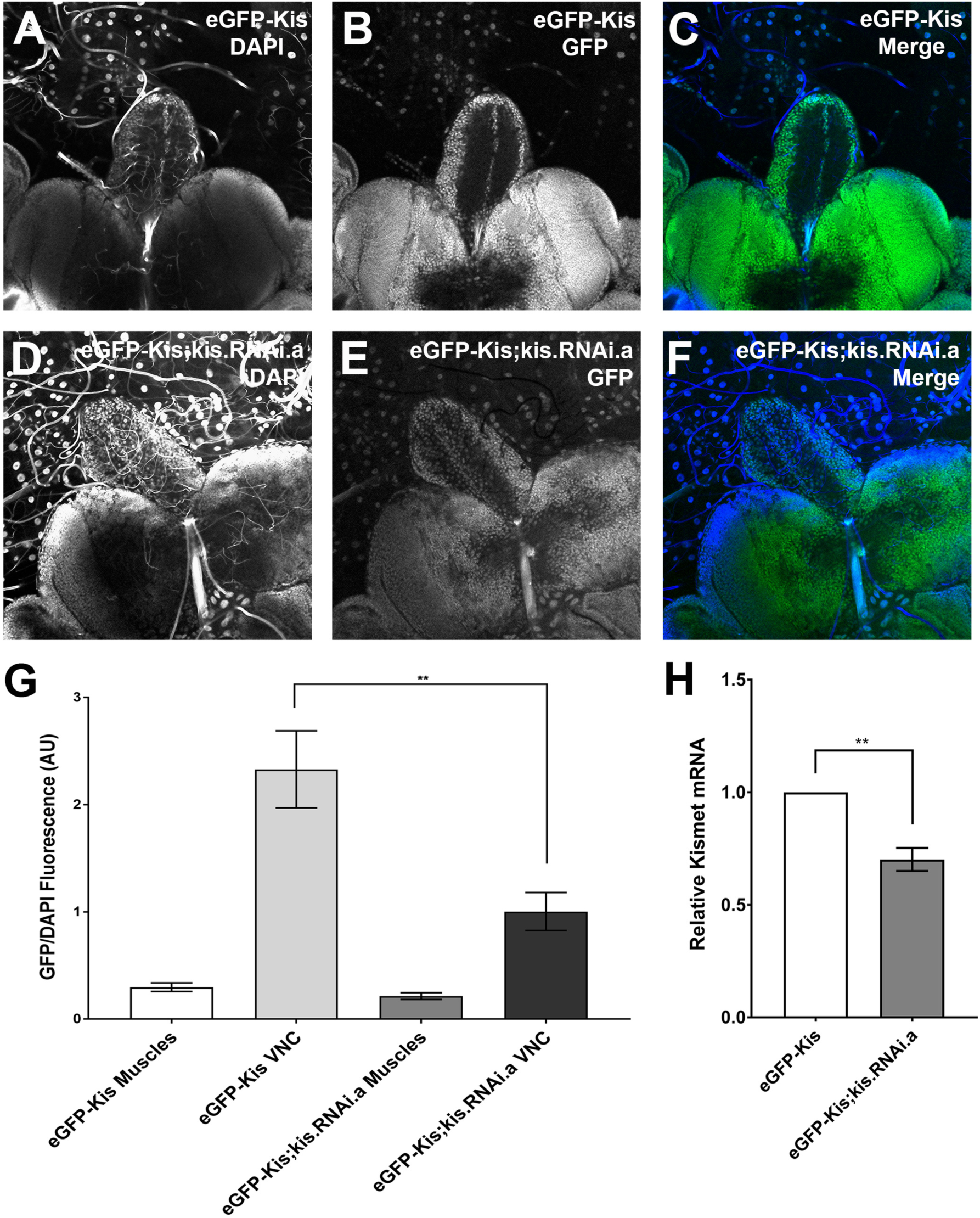
Kis-eGFP is knocked down by Kis.RNAi.a. (A-F) Representative images of Kis-eGFP and *elav*-Gal-4 driven knockdown 3^rd^ instar larval brains stained with DAPI. (G) Quantification of GFP fluorescence intensity compared to that of DAPI (n = 9-12). (H) *Kis* mRNA levels in 3^rd^ instar larval brains of *elav*-Gal-4 driven control and knockdown analyzed by RT-qPCR (n = 4).

**Figure S3.**
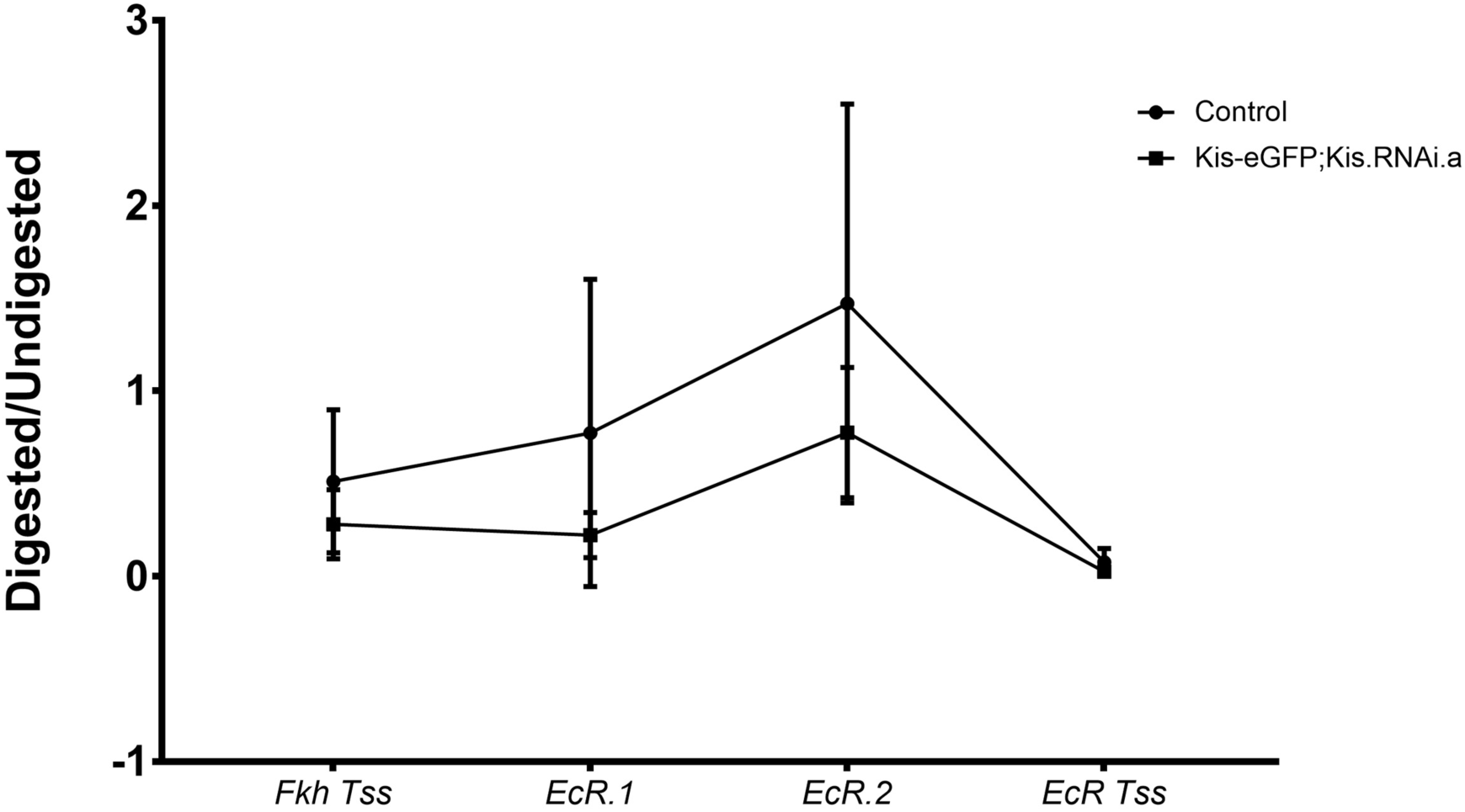
Kis does not alter DNA protection at *EcR* loci. Quantification of MNase protection assay followed by qPCR at the *EcR* TSS, EcR enhancer sites, and the *fkh* TSS (n = 3).

**Figure S4.**
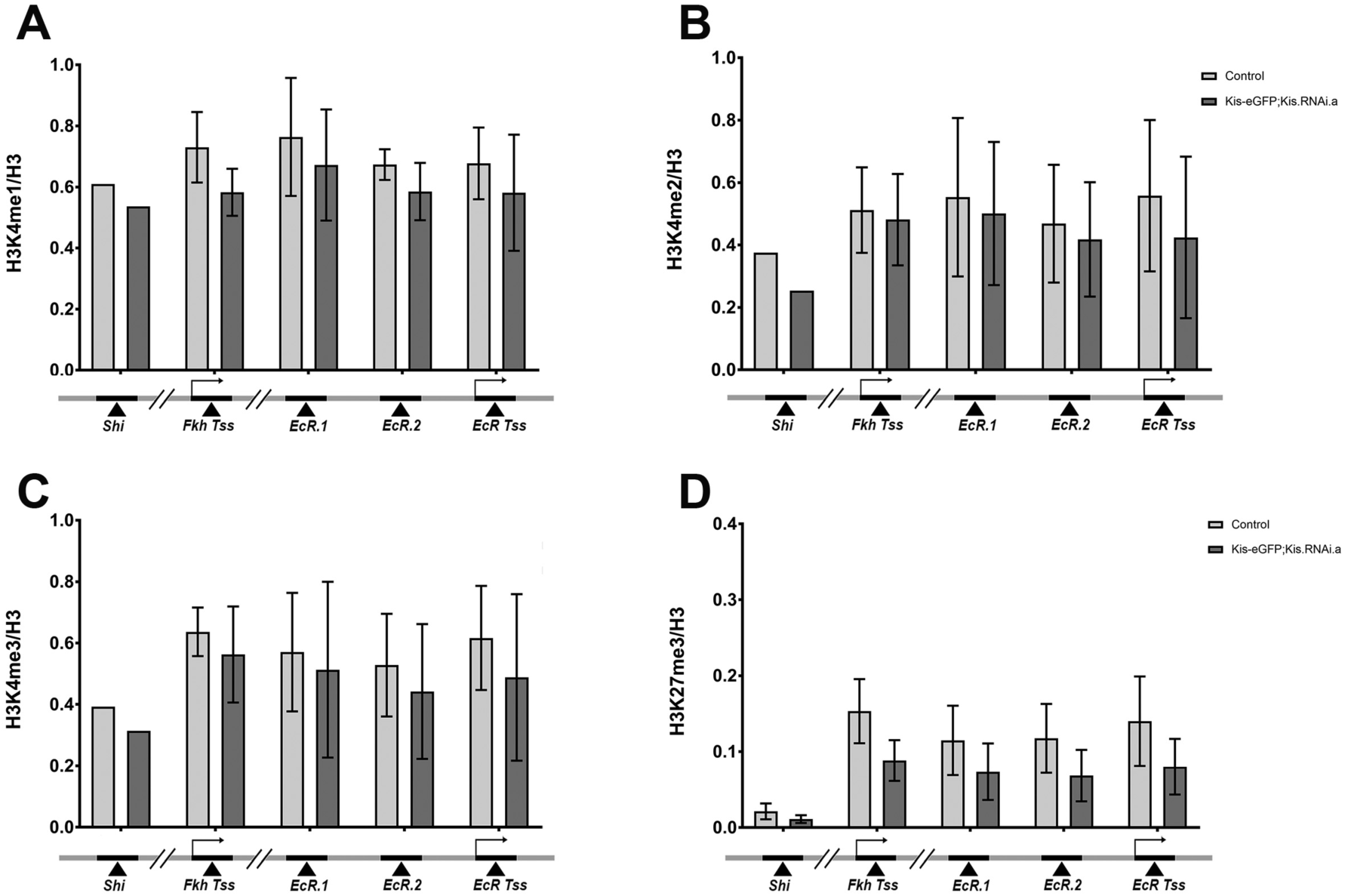
Kis does not alter H3K4 or H3K27 methylation at *EcR* loci. (A-D) qPCR analysis H3K4me1, H3K4me2, H3K4me3, H3K27me3 abundance at the *EcR* TSS and enhancer sites, the *fkh* TSS, and the *shi* promoter site, respectively (n = 2-4).

**Figure S5.**
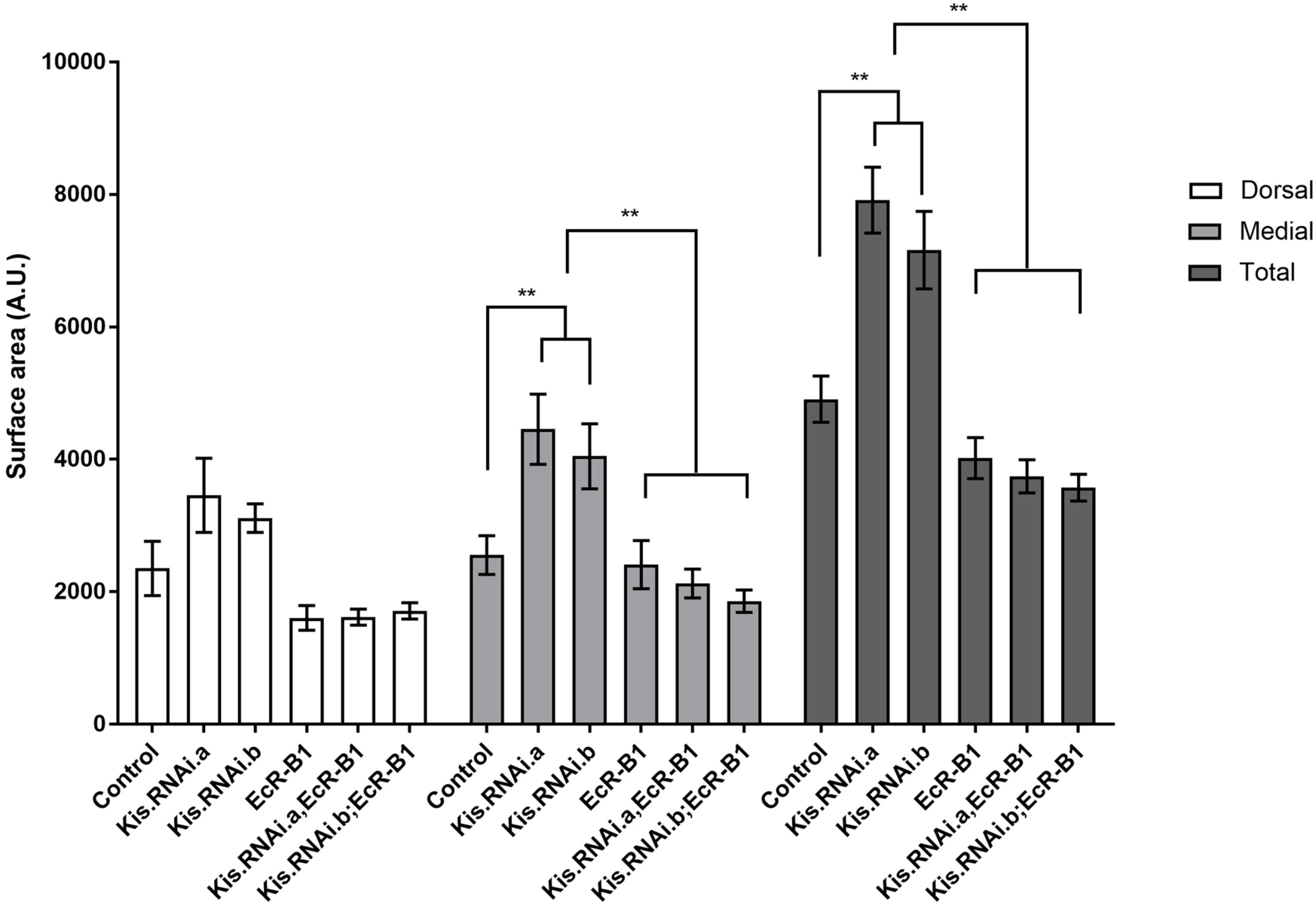
Exogenous EcR-B1 rescues defective axon pruning in Kis knockdown pupae. Quantification of dorsal, medial, and total MB lobe surface areas in e*lav*-Gal-4, mCD8-GFP driven pupae (n = 10).

## Experimental Procedures

### Drosophila stocks and genetics

Unless otherwise noted, all crosses were carried out at 25 °C in a 12:12 light:dark cycle at 60% humidity on standard cornmeal-molasses-agar medium. BL numbers refer to Bloomington Stock Center (BSC) stock numbers (http://flystocks.bio.indiana.edu/bloomhome.htm). VDRC numbers refer to the Vienna Drosophila Resource Center stock numbers (http://stockcenter.vdrc.at/control/main). To drive the expression of transgenes in Drosophila, the Gal-4/UAS bipartite system was used as previously described (Brand and Perrimon, 1993). All Gal-4 stocks (*elav*-Gal-4 (BL #458), *elav*-Gal-4, mCD8-GFP (BL #5146), (*OK107*-Gal-4 (BL #44407)), w^1118^ (BL #5905), UAS:EcRB1 (BL #6469), and Canton S (wild-type) were obtained from Bloomington Stock Center. UAS:Kis.RNAi.a and UAS:Kis.RNAi.b (VDRC #10762 and #46685, respectively) came from the VDRC and were previously described (Melicharek et al., 2010). The *kis*^*LM27*^ allele was generated by EMS mutagenesis, as previously described (Melicharek et al., 2008). UAS:Kis-L, Kis-eGFP, *201Y*-Gal-4 MARCM stocks were gifts from J. Tamkun, A. Spralding, and L. Luo, respectively (Buszczak et al., 2007; Ghosh et al., 2014). MARCM mosaic analysis clones were generated as previously described (Lee and Luo, 1999; Wu and Luo, 2006). The EcR transcriptional reporter (*EcR.1*-Gal-4) was obtained as described (Pfeiffer et al., 2011; Pfeiffer et al., 2008).

Pharmacological treatment media was prepared as described (Latcheva et al., 2018). Treated fly media was made using dried instant food (Nutri-Fly Instant, Genesee Scientific) with water containing 1.6% of 10% w/v tegosept (methyl p-hydroxybenzoate in 95% ethanol) and 0.1% of DMSO vehicle or 10µM SAHA. *Drosophila* were raised on drug containing food for their entire lifespan.

### Immunohistochemistry

Immunohistochemical staining was carried out as previously described (Ghosh et al., 2014). Dissections were performed on Sylgard-coated plates in phosphate buffer and fixed in 2% paraformaldehyde. Primary antibodies obtained from the Iowa Developmental Hybridoma Bank included α-EcR-B1 (1:200), α-Gal-4 (1:200), and α-FasII (1:200). Fluorescently conjugated goat α-rabbit or goat α-mouse secondary antibodies (1:100, Jackson Immunoresearch Labs). Brains were mounted in Vectashield with DAPI (Vector Laboratories, H-1000) and images were obtained using an Olympus Fluoview 1000 laser scanning confocal microscope.

### Quantitative RT-PCR

Pupal (approximately 18 hours APF) heads or third instar larval brains were dissected in ice-cold phosphate buffer and immediately transferred to RNA Later (Abion) and stored in –80 °C. Isolation of total RNA was done using phenol:chloroform extraction followed by alcohol precipitation for purification. RNA was stored in DEPC water at –80 °C. An adapted version of iTaq^™^ Universal SYBR^®^ Green One-Step protocol (Bio-Rad) was utilized and samples were run on Bio-Rad C1000 Thermal Cycler CFX96 Real-Time system. Primers were made to *Kismet, EcR-B1, Gal-4* mRNAs (IDT). ΔC(t) values were calculated by subtracting the C(t) value of each primer set from C(t) value of *RP49* housekeeping control. Fold change in expression was calculated from ΔΔC(t) values. Each experiment was performed in triplicate with at least three biological replicates.

### Chromatin Immunoprecipitation

Brains from third instar larvae were isolated in ice-cold phosphate buffer. Brains were transferred to 1X phosphate buffered saline (PBS, Hyclone) and stored at –80 °C. A modified version of truChIP Tissue Chromatin Shearing Kit with SDS Shearing Buffer protocol (Covaris) was used to shear the DNA. Heads were washed twice with 1X PBS and then fixed in Buffer A with 1% methanol-free formaldehyde for 5 minutes at room temperature. Fixing was stopped with Quenching Buffer E followed by incubation for 5 minutes at room temperature. Tissue was pelleted by centrifugation at 4 °C for 5 minutes. Supernatant was removed, and tissue was washed twice with cold 1X PBS. Wash buffer (WB) was removed and tissue was homogenized for 2-3 minutes in 500µL Lysis Buffer (LB) B. Volume was increased to 1mL with LB B followed by incubation on rocker at 4 °C for 20 minutes with 3 second vortex every 10 minutes. Lysed tissue was pelleted and resuspened in WB C. Tissue was washed on rocker for 10 minutes at 4°C at which time it was pelleted, followed by an additional washing with WB C without incubation. Pelleted lysed and washed tissue, largely consisting of nuclei, was resuspended in Covaris SDS Shearing Buffer D. The aggregate of nuclei was incubated with Buffer D for 10 minutes with occasional vortex prior to transfer to a TC 12X12 tube for shearing. Shearing followed the S- and E-Series Shearing recommendations for 15 minutes. Aliquots were stored at – 80 °C.

Sheared DNA was confirmed to be within a target range of 100-600 bp fragments. Chromatin was immunoprecipitated using Magna ChIP™ HiSens kit (Millipore). Antibodies against modifications H3K27me3 (rabbit, ab195477), H3K4me1 (rabbit, ab8895), H3K4me2 (rabbit, ab7766), H3K4me3 (rabbit, ab8580), H3K36me2 (rabbit, ab9049), H3K36me3 (rabbit, ab9050), H4K16ac (rabbit, emd millipore 07-329) were used and compared to Histone H3 (rabbit, ab1791) and Histone H4 (rabbit, ab10158) antibodies as appropriate. To examine Kismet abundance, α-GFP (rabbit, ab290) was used and α-IgG (rabbit, ab171870) was used as a background control. After elution, samples were incubated with RNaseA (10mg/mL, ThermoScientific) at 37 °C for 30 minutes followed by an incubation with proteinase K (10mg/mL, Millipore) at 57 °C overnight and then inactivate at 75 °C for 15 minutes the next day. Isolated DNA was purified via QIAquick^®^ PCR Purification Kit (Qiagen) and stored at –20 °C.

### MNase Protection Assay

Brains from third instar larvae were dissected in ice-cold phosphate buffer, transferred to 1X PBS, and stored at –80 °C. An MNase protection assay was performed using an adapted protocol from (Berson et al., 2017; Chereji et al., 2016). Tissue was homogenized in 500µL of crosslinking buffer (60mM KCl, 15mM NaCl, 4mM MgCl2,15mM HEPES pH 7.6, 0.5mM DTT, 0.5% Triton X-100, protease inhibitor (100X), 2% formaldehyde) and incubated at room temperature for 15 minutes. Crosslinking was quenched with 50µL of 2.5M of glycine and incubated at room temperature for 5 minutes. Samples were washed twice in crosslinking buffer and twice in D1 buffer (25% glycerol, 5mM Mg Acetate, 50mM Tris pH 8.0, 0.1mM EDTA, 5mM DTT) and resuspended in 1mL of MNase buffer (60mM KCl, 15mM NaCl, 15mM Tris pH 7.4, 0.5mM DTT, 0.25M sucrose, 1.0mM CaCl2). 10units MNase (70196Y, Affymetrix) was added to sample tubes and incubated at 37 °C for 30 minutes. Reaction was quenched with EDTA (final 12.5mM) and SDS (final 0.5%). Samples were equilibrated with NaCl (final 140mM) and incubated with RNaseA (10mg/mL) at 37 °C and then overnight with proteinase K (10mg/mL) at 65 °C and then 15 minutes at 75 °C the next day. DNA was purified via QIAquick PCR Purification Kit and stored at –20 °C.

### Q-PCR

Purified DNA was used to prep PCR reaction mixes according to DyNAmo Flash SYBR Green qPCR Kit (ThermoFisher Scientific). Samples were run using Bio-Rad C1000 Thermal Cycler CFX96 Real-Time system. Primers were made to the TSS of *EcR-B1*, a 16kb upstream enhancer of *EcR-B1* (*EcR.1*), and a 7.4kb upstream enhancer of *EcR-B1* (*EcR.2*) (IDT). Additionally, positive and negative control primers were made to the TSS of *fkh* and the *shi* promoter site, respectively (IDT). For control and RNAi knockdown analysis values were adjusted for input at each primer set and ΔΔC(t) values were calculated by subtracting the adjusted ΔC(t) value of each primer set from the corresponding ΔC(t) IgG control. Fold change in expression was calculated from ΔΔC(t) values. Each experiment was performed in triplicate with at least three biological replicates.

### Behavioral testing

To evaluate learning and memory, the canonical fly courtship behavior was used as a readout in an associative conditioning assay described by Siegel and Hall (Siegel and Hall, 1979). Virgin male flies (0 to 6 hours following eclosion) were collected in individual food vials and aged 5 days. Similarly, virgin female wild-type flies were collected, transferred to collective food vials, and aged 5 days. 24 hours before assessment, virgin wild-type females were mated individually using wild-type males. These flies were subsequently separated from virgin females. This behavioral test was executed in a separate room kept at 25°C and 50% humidity, recorded using a Sony DCR-SR47 Handycam with Carl Zeiss optics, and illuminated from below using a constant 115V white light transilluminator. Genotypes of each male were blinded on the day of the assay and all fly transfers were performed without anesthesia. Aged male flies were transferred to mating chambers (Aktogen) each containing a portioned-off mated female fly. Flies were allowed to acclimate for 2 minutes before the assay. Training was recorded and commenced for 60 minutes. After, the male fly was transferred to a clean mating chamber containing a portioned-off virgin female fly. After an additional 2-minute acclimation period, the divider was removed, and immediate recall was recorded for 10 minutes. Shams experienced the same manipulations however these aged males were not exposed to any fly during the training portion. Digital video analysis of the time spent courting was performed using iMovie software (Apple). Courtship indices were calculated by total time observed performing courtship behaviors divided by total time assayed.

### Statistical analysis

All statistical analyses were performed using GraphPad Prism (v. 7.03). Significance was determined at the 95% confidence interval. Unpaired student’s *t*-test was used for all experiments, except pupal pruning analysis (utilized one-way ANOVA test) and the learning portion of the associative conditioning assay (utilized paired student’s *t*-test). Statistical significance in figures is represented by * = p < 0.05, ** = p < 0.001, *** = p < 0.0001. Error bars represent the standard error of the mean (SEM).

